# Deep image reconstruction from human brain activity

**DOI:** 10.1101/240317

**Authors:** Guohua Shen, Tomoyasu Horikawa, Kei Majima, Yukiyasu Kamitani

**Author notes:** These authors contributed equally to this work (Guohua Shen, Tomoyasu Horikawa, Kei Majima).

## Abstract

Machine learning-based analysis of human functional magnetic resonance imaging (fMRI) patterns has enabled the visualization of perceptual content. However, it has been limited to the reconstruction with low-level image bases (Miyawaki et al., 2008; Wen et al., 2016) or to the matching to exemplars (Naselaris et al., 2009; Nishimoto et al., 2011). Recent work showed that visual cortical activity can be decoded (translated) into hierarchical features of a deep neural network (DNN) for the same input image, providing a way to make use of the information from hierarchical visual features (Horikawa & Kamitani, 2017). Here, we present a novel image reconstruction method, in which the pixel values of an image are optimized to make its DNN features similar to those decoded from human brain activity at multiple layers. We found that the generated images resembled the stimulus images (both natural images and artificial shapes) and the subjective visual content during imagery. While our model was solely trained with natural images, our method successfully generalized the reconstruction to artificial shapes, indicating that our model indeed ‘reconstructs’ or ‘generates’ images from brain activity, not simply matches to exemplars. A natural image prior introduced by another deep neural network effectively rendered semantically meaningful details to reconstructions by constraining reconstructed images to be similar to natural images. Furthermore, human judgment of reconstructions suggests the effectiveness of combining multiple DNN layers to enhance visual quality of generated images. The results suggest that hierarchical visual information in the brain can be effectively combined to reconstruct perceptual and subjective images.

---

Whereas it has long been thought that the externalization or visualization of states of the mind is a challenging goal in neuroscience, brain decoding using machine learning analysis of fMRI activity nowadays has enabled the visualization of perceptual content. Although sophisticated decoding and encoding models have been developed to render human brain activity into images or movies, the methods were essentially limited to the image reconstruction with low-level image bases (Miyawaki et al., 2008; Wen et al., 2016) or to the matching to exemplar images or movies (Naselaris et al., 2009; Nishimoto et al., 2011), failing to combine visual features of multiple hierarchical levels. Furthermore, while several recent attempts introduced deep neural networks (DNNs) into visual image reconstructions, they also did not fully utilize hierarchical information to reconstruct visual images (Seeliger et al., 2017, Han et al., 2017).

The recent success of deep neural networks provides technical innovations to study the hierarchical visual processing in computational neuroscience (Yamins & DiCarlo, 2016). Our recent study used DNN visual features as a proxy for the hierarchical neural representations of the human visual system, and found that a brain activity pattern measured by fMRI can be decoded (translated) into DNN features given the same input (Horikawa & Kamitani, 2017). The finding revealed a homology between the hierarchical representations of the brain and the DNN, providing a new opportunity to make use of the information from hierarchical visual features.

Here, we present a novel approach, named *deep image reconstruction*, to visualize perceptual content from human brain activity. We combined the DNN feature decoding from fMRI signals and the methods for image generation recently developed in the machine learning field (Mahendran & Vedaldi, 2015) (Fig. 1). The reconstruction algorithm starts from a random image and iteratively optimize the pixel values so that the DNN features of the input image become similar to those decoded from brain activity across multiple DNN layers. The resulting optimized image is taken as the reconstruction from the brain activity. We also introduced a deep generator network (DGN) (Nguyen et al., 2016) as a prior to constrain reconstructed images to be similar to natural images.

**Figure 1.**
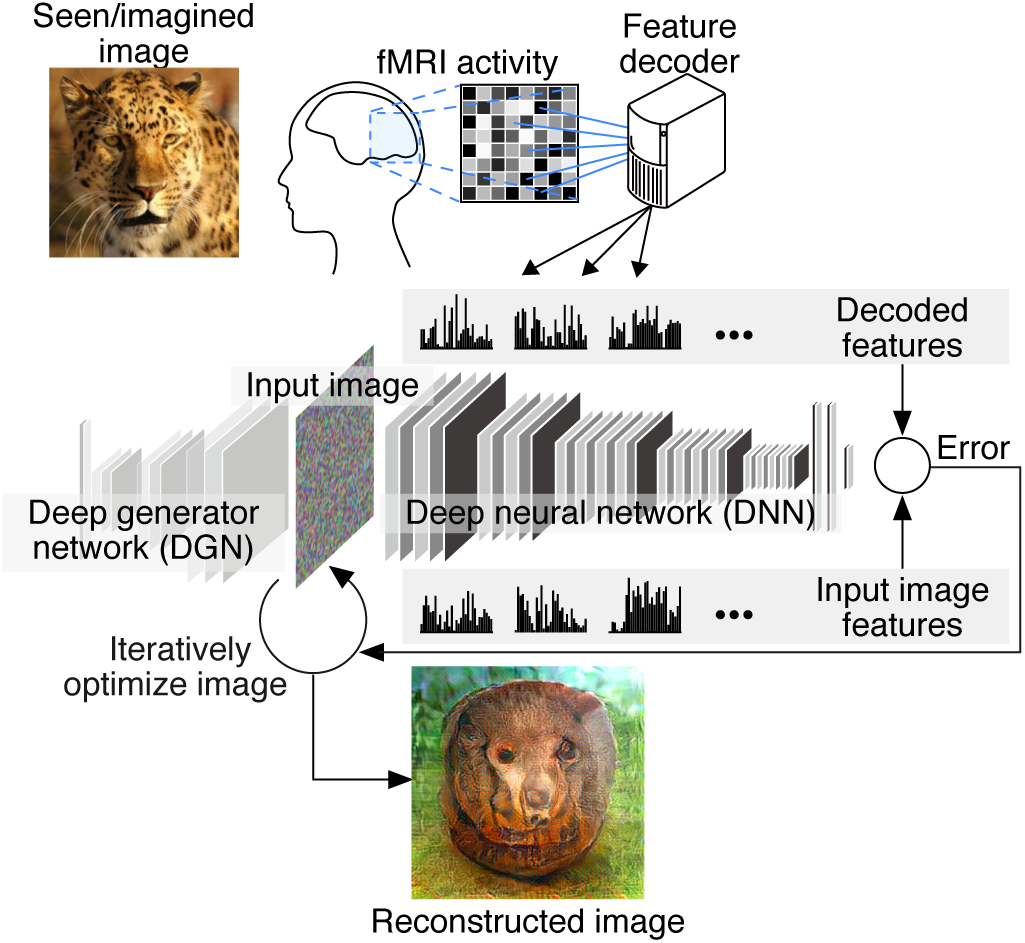
Deep image reconstruction. Overview of deep image reconstruction is shown. The pixels’ values of the input image are optimized so that the DNN features of the image are similar to those decoded from fMRI activity. A deep generator network (DGN) is optionally combined with the DNN to produce natural-looking images, in which optimization is performed at the input space of the DGN.

We trained the decoders that predict the DNN features of viewed images from fMRI activity patterns, following the procedures of Horikawa and Kamitani (2017). Our experiments consisted of four distinct types of image presentation sessions: training natural-image sessions, test natural-image sessions, geometric-shape sessions, and alphabetical-letter sessions, and one mental-imagery session. The decoders were trained using fMRI data measured while subjects were viewing natural images. The trained decoders were then used to predict DNN features from independent test fMRI data collected during the presentation of novel natural images and artificial shapes and during mental imagery (Supplementary Fig. 1). Then, the decoded features were forwarded to the reconstruction algorithm.

The reconstructions of the natural images from three subjects are shown in Fig. 2a (see Supplementary Fig. 2 for more examples). These reconstructions obtained with the DGN capture the objects’ dominant structures in the images. Furthermore, fine structures reflecting semantic aspects like faces, eyes, and texture patterns were also generated in several images. This is not likely a coincidence because such fine structures were robustly observed through the optimization processes (Supplementary Fig. 3, and Supplementary Movie 1^1^). The robustness of the analyses was also confirmed with a previously published dataset (Horikawa & Kamitani, 2017), reproducing quantitatively similar reconstructions with those in the present study (Supplementary Fig. 4).

**Figure 2.**
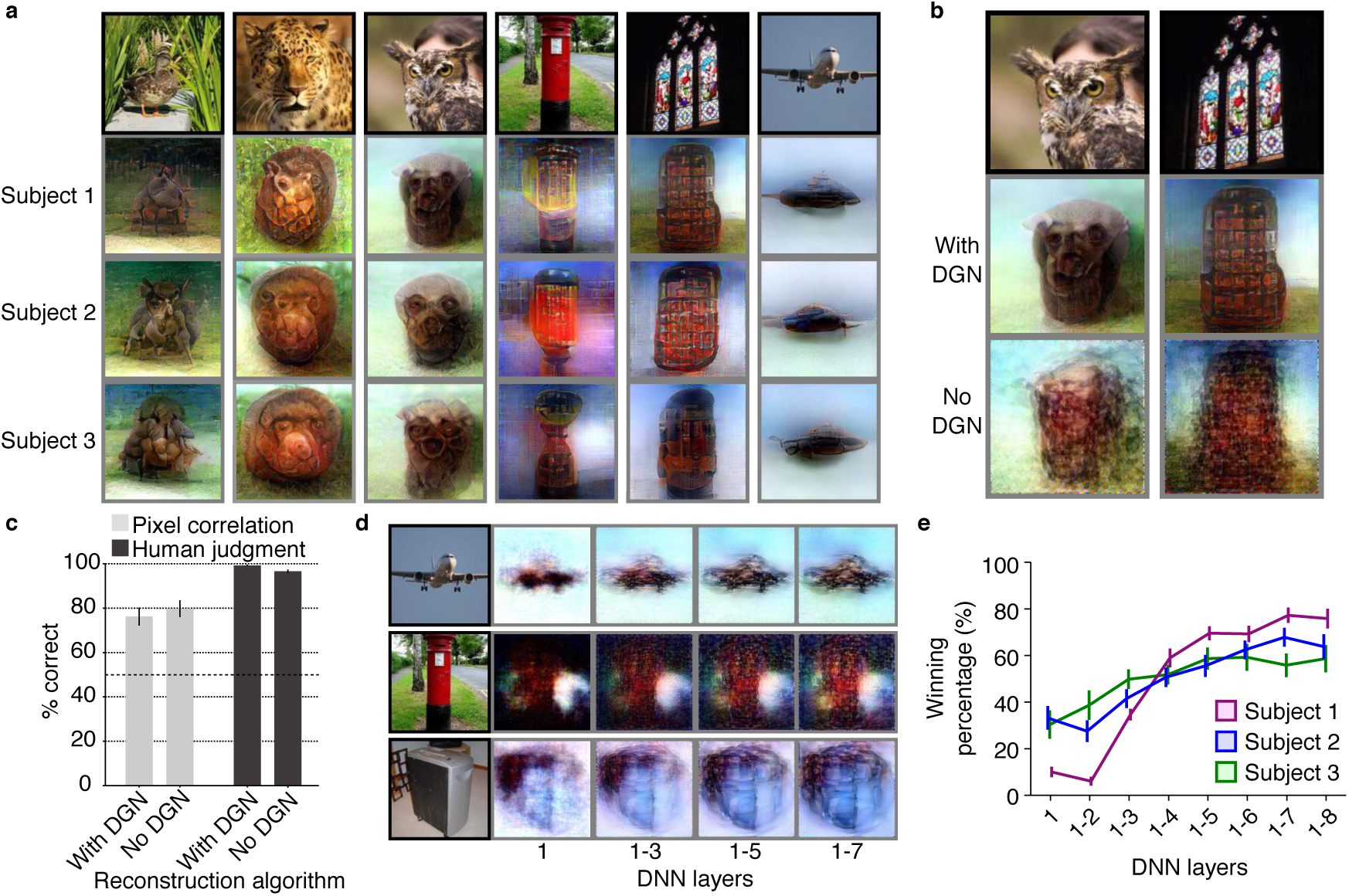
Seen natural image reconstructions. Images with black and gray frames show presented and reconstructed images, respectively (reconstructed from VC activity). **a**, Reconstructions utilizing the DGN (using DNN1–8). Three reconstructed images correspond to reconstructions from three subjects. **b**, Reconstructions with and without the DGN (DNN1–8). The first, second, and third rows show presented images, reconstructions with and without the DGN, respectively, **c**, Reconstruction quality of seen natural images (error bars, 95% confidence interval (C.I.) across samples; three subjects pooled; chance level, 50%). **d**, Reconstructions using different combinations of DNN layers (without the DGN). **e**, Subjective assessment of reconstructions from different combinations of DNN Sayers (errorbars, 95% C.I. across samples).

To investigate the effect of the natural image prior, we compared the reconstructions generated with and without the DGN (Fig. 2b). While the reconstructions obtained without the DGN also successfully reconstructed rough silhouettes of dominant objects, they did not show semantically meaningful appearances (Supplementary Fig. 5). Evaluations by pixel-wise spatial correlation and by human judgment both showed almost comparative accuracy for reconstructions with and without the DGN (by pixel-wise spatial correlation, with and without the DGN, 76.1% and 79.7%; by human judgment, with and without the DGN, 99.1%, 96.5%). However, reconstruction accuracy evaluated by pixel-wise spatial correlation showed slightly higher accuracy with reconstructions without the DGN than those with the DGN (two-sided signed-rank test, *p* < 0.01), whereas the opposite was observed with evaluations by human judgment (two-sided signed-rank test, *p* < 0.01). These results suggest the utility of the DGN that enhances perceptual similarity of reconstructed images to target images by rendering semantically meaningful details to reconstructions.

To characterize the “deep” nature of our method, the effectiveness of combining multiple DNN layers was tested using human rating. For each of the 50 test natural images, reconstructed images were generated with a variable number of multiple layers (Fig. 2d, and Supplementary Fig. 6; DNN1 only, DNN1–2, DNN1–3, …, DNN1–8). An independent rater was presented with an original image and a pair of reconstructed images, both from the same original image but generated with different combinations of multiple layers, and indicated which of the reconstructed images looked more similar to the original image. The subjective assessment showed higher rates of choice with reconstructions from a larger number of DNN layers (Fig. 2e), suggesting the improvement of reconstruction quality by combining multiple levels of visual features.

To confirm that our method was not restricted within the specific image domain used for the model training, we tested whether it is possible to generalize the reconstruction to artificial shapes. This was challenging because both the DNN and our decoding models were solely trained on natural images. The reconstructions for colored artificial shapes are shown in Fig. 3a (also see Supplementary Fig. 7 for more examples). The results show that artificial shapes were successfully reconstructed with moderate accuracy (Fig. 3b; 69.4% by pixel-wise spatial correlation, 92.3% by human judgment), indicating that our model indeed ‘reconstructs’ or ‘generates’ images from brain activity, not simply matches to exemplars.

**Figure 3.**
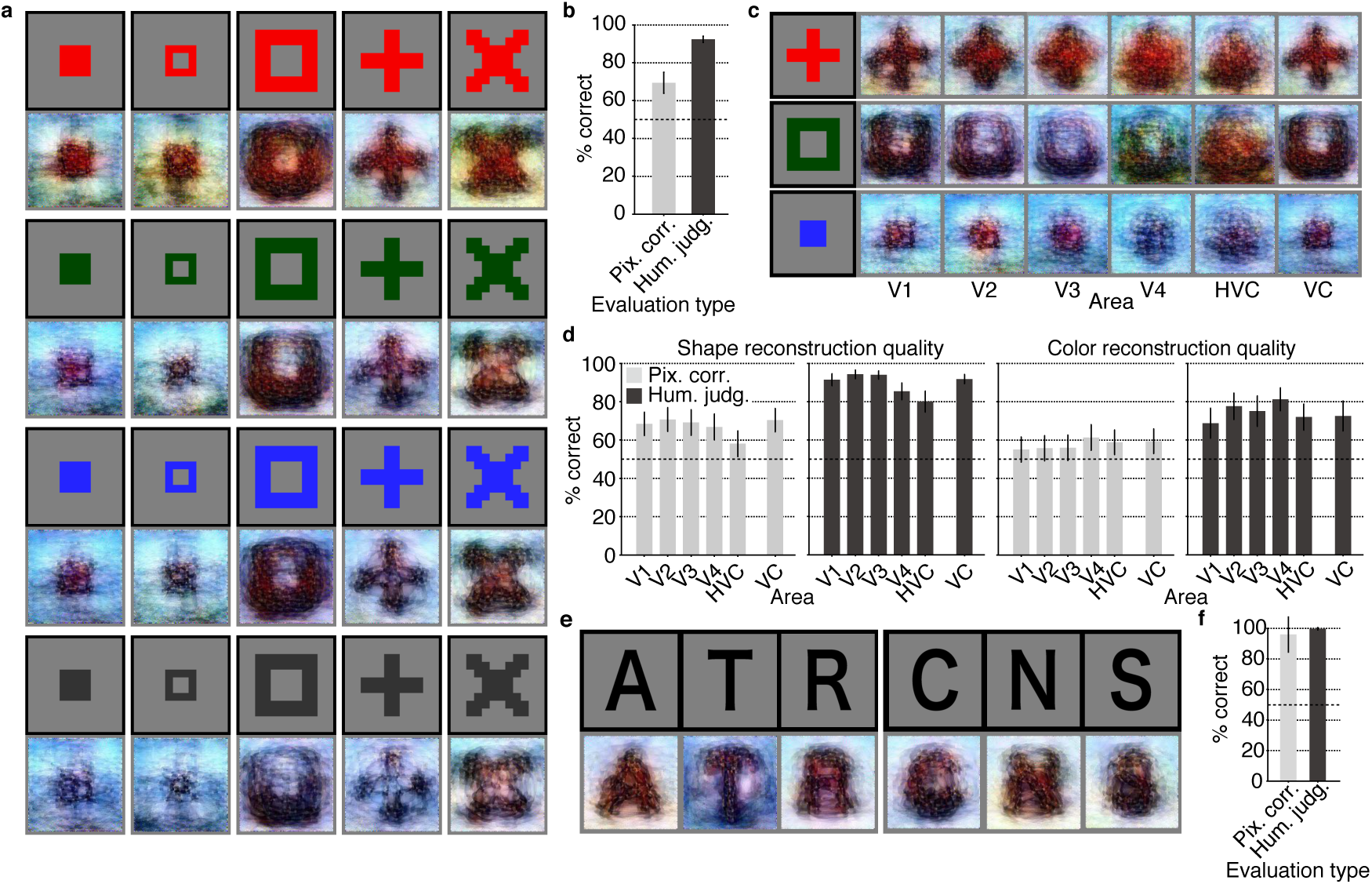
Seen artificial shape reconstructions. Images with black and gray frames show presented and reconstructed images (DNN 1∓8, without the DGN). **a**, Reconstructions for seen colored artificial shapes (VC activity). **b**, Reconstruction quality of colored artificial shapes. **c**, Reconstructions of colored artificial shapes obtained from multiple visual areas. **d**, Reconstruction quality of shape and colors for different visual areas, **f**, Reconstructions of alphabetical letters. **f**, Reconstruction quality for alphabetical letters. For **b**, **d**, **f**, error bars indicate 95% C.I. across samples (three subjects pooled; chance level, 50%).

To assess how the shapes and colors of the stimulus images were reconstructed, we separately evaluated the reconstruction quality of each of shape and color by comparing reconstructed images of the same colors and shapes. Analyses with different visual areas show different tendency of reconstruction quality for shapes and colors (Fig. 3c and Supplementary Fig. 8). Evaluations by human judgment suggest that shapes were better reconstructed from early visual areas, whereas colors were better reconstructed from mid-level visual areas, V4 (Fig. 3d; ANOVA, interaction, between task type (shape vs. color) and brain areas, *p* < 0.05), while the interaction effect was marginal from evaluations by pixel-wise spatial correlation *(p* > 0.05). These contrasting patterns further support the success of shape and color reconstructions, and indicate that our method can be a useful tool to characterize information content encoded in activity patterns in individual brain areas by visualization.

We also tested reconstructions of alphabetical letters to obtain a performance benchmark (Fig. 3e, and Supplementary Fig. 9). The reconstructed letters were arranged in a word: “ATR” and “CNS”. The evaluations of reconstruction quality showed high accuracy for these letters (Fig. 3f; 95.9% by pixel-wise spatial correlation, 99.6% by human judgment). The successful reconstructions of alphabetical letters demonstrate that our method expands possible states of visualizations by advancing the resolution of reconstructions from previous studies (Miyawaki et al., 2008; Schoenmakers et al., 2013).

Finally, to explore the possibility of visually reconstructing subjective content, we performed an experiment in which participants were asked to produce mental imagery of natural images and artificial images shown prior to the task session. The reconstructions from brain activity due to mental imagery are shown in Fig. 4 (see Supplementary Fig.10 for more examples). While reconstruction quality varied across subjects and images, rudimentary reconstructions were obtained for some of the artificial shapes (Fig. 4a). In contrast, imagined natural images were not well reconstructed, possibly because of the difficulty to imagine complex natural images (Fig. 4b). While evaluations of reconstructed artificial shapes by pixel-wise spatial correlation failed to show high accuracy (Fig. 4c; 51.9%), this may be due to the possible disagreements of positions, colors and luminance between target and reconstructed images. Meanwhile, evaluations by human judgment showed higher than the chance accuracy, suggesting that imagined shapes were recognizable from reconstructed images (Fig. 4c; 83.2%). Furthermore, separate evaluations for color and shape reconstructions of artificial images suggest that shape rather than color highly contributed to the high proportion of correct answers by human raters (Fig. 4d; color, 64.8%; shape, 87.0%; two-sided signed-rank test, *p* < 0.01). Taken together, these results provide evidence for the feasibility of visualizing imagined content from brain activity patterns.

**Figure 4.**
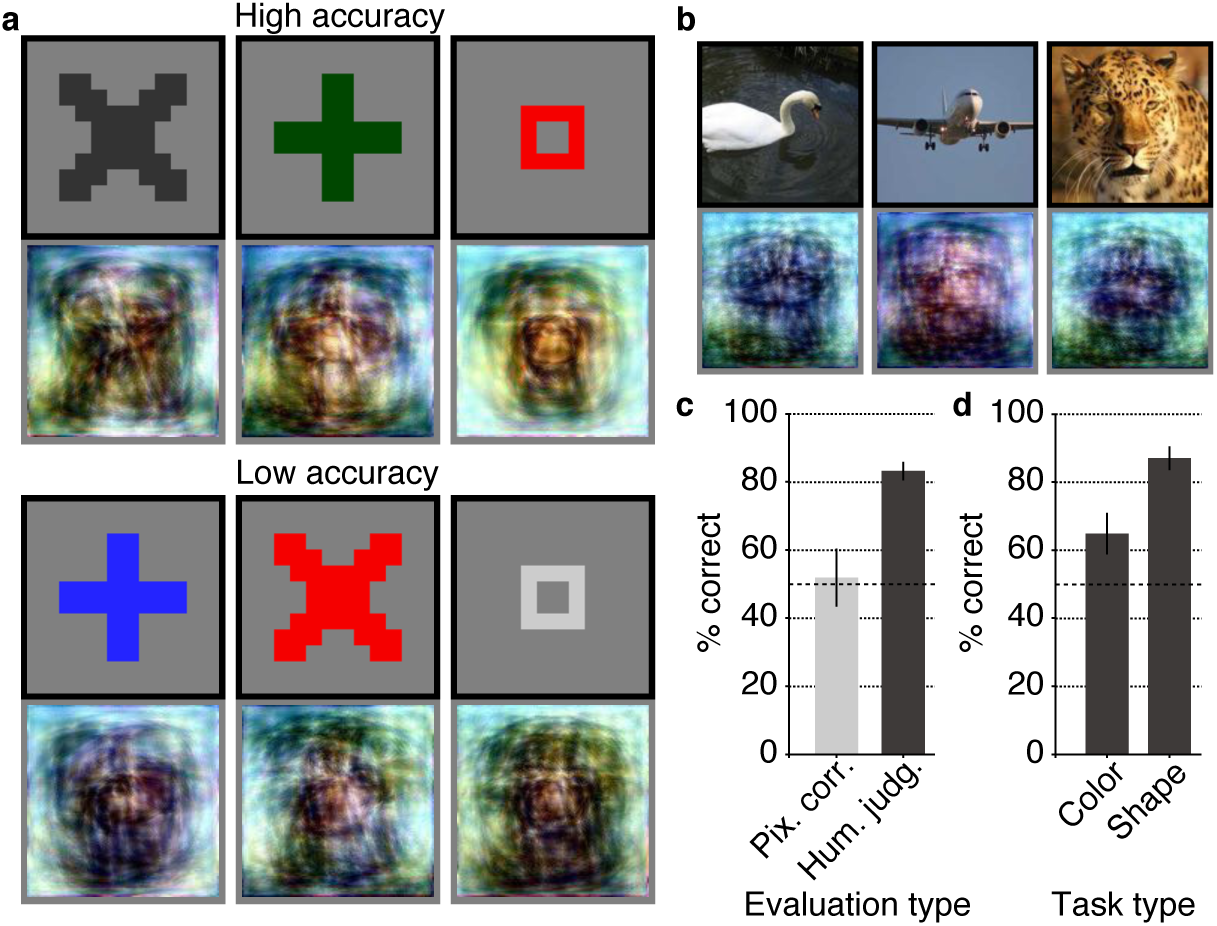
Imagery reconstructions. Images with black and gray frames show target and reconstructed images (VC activity, DNN 1–8, without the DGN). **a**, Reconstructions for imagined artificial shapes. Images with high/low accuracy evaluated by human judgment are shown. **b**, Reconstructions for imagined natural images. **c**, Reconstruction quality of imagery reconstructions (error bars,95% C.I. across samples; three subjects pooled; chance level, 50%). **d**, Reconstruction quality of artificial shapes separately evaluated for color and shape by human judgment (error bars, 95% C.I, across samples; three subjects pooled; chance level, 50%).

We have presented a novel approach to reconstruct perceptual and mental content from human brain activity combining visual features from the multiple layers of a DNN. We successfully reconstructed viewed natural images, especially when combined with a DGN. Reconstruction of artificial shapes was also successful, even though the reconstruction models used were trained only on natural images. The same method was applied to imagery to reveal rudimentary reconstructions of mental content. Our approach could provide a unique window into our internal world by translating brain activity into images via hierarchical visual features.

One interesting observation that can be obtained from the present results is that the luminance contrast of a reconstruction was often reversed (e.g., stained glass images in Fig. 2a), presumably because of the lack of (absolute) luminance information in fMRI signals even in the early visual areas (Haynes et al., 2004). Additional analyses revealed that feature values of filters with high luminance contrast in the earliest DNN layers (conv1_1 in VGG19) were better decoded when they were converted to absolute values (Supplementary Figure 11), demonstrating a clear discrepancy of fMRI and raw DNN signals. While this may limit potential for reconstructions from fMRI signals, the ambiguity might be resolved by modeling DNNs to fill gaps between DNNs and fMRI. Alternatively, emphasizing high-level visual information in hierarchical visual features may help to resolve the ambiguity of luminance by incorporating information about semantic context.

## Methods

### Subjects

Three healthy subjects with normal or corrected-to-normal vision participated in our experiments: Subject 1 (male, age 33), Subject 2 (male, age 23) and Subject 3 (female age 23). This sample size was chosen based on previous fMRI studies with similar experimental designs (Miyawaki et al., 2008; Horikawa & Kamitani, 2017). All subjects provided written informed consent for participation in the experiments, and the study protocol was approved by the Ethics Committee of ATR.

### Visual stimuli

Visual stimuli consisted of natural images, artificial geometric shapes, and alphabetical letters. The natural images were identical to those used in Horikawa & Kamitani (2017), which were originally collected from an online image database *ImageNet* (Deng et al., 2009; 2011, fall release). Those images were cropped to the center and were resized to 500 × 500 pixels. The artificial geometric shapes consisted of a total of 40 combinations of 5 shapes and 8 colors (red, green, blue, cyan, magenta, yellow, white, and black), in which the shapes were identical to those used in Miyawaki et al. (2008) and the luminance were matched across colors except for white and black. The alphabetical letter images consisted of 10 black letters, including “A”, “C”, “E”, “I”, “N”, “O”, “R”, “S”, “T”, and “U”.

### Experimental design

We conducted two types of experiments: image presentation experiments and an imagery experiment. The image presentation experiments consisted of four distinct session types, in which different variants of visual images were presented (natural images, geometric shapes, and alphabetical letters). All visual stimuli were rear-projected onto a screen in an fMRI scanner bore using a luminance-calibrated liquid crystal display projector. To minimize head movements during fMRI scanning, subjects were required to fix their heads using a custom-molded bite-bar made for each subject. Data from each subject were collected over multiple scanning sessions spanning approximately 10 months. On each experiment day, one consecutive session was conducted for a maximum of 2 hours. Subjects were given adequate time for rest between runs (every 5–8 min) and were allowed to take a break or stop the experiment at any time.

### Image presentation experiment

The image presentation experiments consisted of four distinct types of sessions: training natural-image sessions, test natural image sessions, geometric-shape sessions, and alphabetical-letter sessions. Each session consisted of 24, 24, 20, and 12 separate runs, respectively. For those four sessions, each run comprised 55, 55, 44, and 11 stimulus blocks consisting of 50, 50, 40, and 10 blocks with different images and 5, 5, 4, and 1 randomly interspersed repetition blocks where the same image as in the previous block was presented (7 min 58 s for the training and test natural-image sessions, 6 min 30 s for the geometric-shape sessions, and 5 min 2 s for the alphabetical-letter sessions, for each run). Each stimulus block was 8 s (training natural-images, test natural-images, and geometric-shapes) or 12 s (alphabetical-letters) long followed by either no (training natural-images, test natural-images, and geometric-shapes) or a 12 s (alphabetical-letters) rest period. Images were presented at the center of the display with a central fixation spot and were flashed at 2 Hz (12 × 12 and 0.3 × 0.3 degrees of visual angle for visual images and fixation spot, respectively). The color of the fixation spot changed from white to red for 0.5 s before each stimulus block began to indicate the onset of the block. Additional 32- and 6-s rest periods were added to the beginning and end of each run, respectively. Subjects were requested to maintain steady fixation throughout each run and performed a one-back repetition detection task on the images, responding with a button press for each repetition to maintain their attention on the presented images (mean task performance across three subjects; sensitivity 0.9820; specificity 0.9995; pooled across sessions). In one set of training natural-image session, a total of 1,200 images were presented only once. This set of training natural-image session was repeated five times. In the test natural-image session, the geometric-shape session, and the alphabetical-letter session, 50, 40, and 10 images were presented 24, 20, and 12 times each, respectively. The presentation order of the images was randomized across runs.

### Imagery experiment

In the imagery experiment, subjects were required to visually imagine (remember) one of 25 images that were selected from those presented in the test natural image session and the geometric shape session of the image presentation experiment (10 natural images and 15 geometric shapes). Prior to the experiment, subjects were asked to relate words and visual images so that they can remember visual images from word cues. The imagery experiment consisted of 20 separate runs and each run contained 26 blocks (7 min 34 s for each run). The 26 blocks consisted of 25 imagery trials and a fixation trial, in which subjects were required to maintained steady fixation without any imagery. Each imagery block consisted of a 4-s cue period, an 8-s imagery period, a 3-s evaluation period and a 1-s rest period. Additional 32- and 6-s rest periods were added to the beginning and end of each run, respectively. During the rest periods, a white fixation spot was presented at the center of the display. The color of the fixation spot changed from white to red for 0.5 s to indicate the onset of the blocks from 0.8 s before each cue period began. During the cue period, words specifying the visual images to be imagined were visually presented around the center of the display (1 target and 25 distractors). The position of each word was randomly changed across blocks to avoid contamination of cue-specific effects on the fMRI response during imagery periods. The word corresponding to the image to be imagined was presented in red (target) and the other words were presented in black (distractors). Subjects were required to start imagining a target image immediately after cue words disappeared. The imagery period was followed by a 3-s evaluation period, in which the word corresponding to the target image and a scale bar was presented to allow the subjects to evaluate the correctness and vividness of their mental imagery on a five-point scale (very vivid, fairly vivid, rather vivid, not vivid, cannot correctly recognize the target) by pressing the left and right buttons of a button box that was placed in their right hand to change the score from its random initial score. As an aid for remembering the associations between words and images, subjects were able to see word and visual image pairs during every inter-run-rest period by controlling buttons so they can confirm their own memory.

### MRI acquisition

fMRI data were collected using 3.0-Tesla Siemens MAGNETOM Verio scanner located at the Kokoro Research Center, Kyoto University. An interleaved T2*-weighted gradient-EPI scan was performed to acquire functional images covering the entire brain (TR, 2,000 ms; TE, 43 ms; flip angle, 80 deg; FOV, 192 **×** 192 mm; voxel size, 2 **×** 2 **×** 2 mm; slice gap, 0 mm; number of slices, 76; multiband factor, 4). T2-weighted turbo spin echo images were scanned to acquire high-resolution anatomical images of the same slices used for the EPI (TR, 11,000 ms; TE, 59 ms; flip angle, 160 deg; FOV, 192 **×** 192 mm; voxel size, 0.75 **×** 0.75 **×** 2.0 mm). T1-weighted magnetization-prepared rapid acquisition gradient-echo (MP-RAGE) fine-structural images of the entire head were also acquired (TR, 2,250 ms; TE, 3.06 ms; TI, 900 ms; flip angle, 9 deg, FOV, 256 **×** 256 mm; voxel size, 1.0 **×** 1.0 **×** 1.0 mm).

### MRI data preprocessing

The first 8-second scans of each run were discarded to avoid MRI scanner instability. We then subjected the acquired fMRI data to three-dimensional motion correction with SPM5 (http://www.fil.ion.ucl.ac.uk/spm). Those data were then coregistered to the within-session high-resolution anatomical images of the same slices used for EPI and subsequently to the whole-head high-resolution anatomical images. The coregistered data were then re-interpolated as 2 × 2 × 2 mm voxels.

Data samples were created by first regressing out nuisance parameters, including a linear trend, and temporal components proportional to six motion parameters calculated by the SPM5 motion correction procedure, from each voxel amplitude for each run. After that, voxel amplitudes were normalized relative to the mean amplitude of the initial 24-s rest period of each run, and were *despiked* to reduce extreme values (beyond ±3SD for each run). The voxel amplitudes were then averaged within each 8-s (training natural image-sessions), 12-s (test natural-image, geometric-shape, and alphabetical-letter sessions) stimulus block (four or six volumes) or 16-s imagery block (eight volumes, imagery experiment), after shifting the data by 4 s (two volumes) to compensate for hemodynamic delays.

### Region of interest (ROI)

V1, V2, V3, and V4 were delineated by the standard retinotopy experiment (Engel et al., 1994; Sereno et al., 1995). The lateral occipital complex (LOC), the fusiform face area (FFA), and the parahippocampal place area (PPA) were identified using conventional functional localizers (Kanwisher et al., 1997; Kourtzi and Kanwisher, 2000; Epstein & Kanwisher, 1998). See Supplementary Methods for details. A contiguous region covering LOC, FFA, and PPA was manually delineated on the flattened cortical surfaces, and the region was defined as the “higher visual cortex” (HVC). Voxels overlapping with V1-V3 were excluded from the HVC. Voxels from V1–V4 and the HVC were combined to define the “visual cortex” (VC). In the regression analysis, voxels showing the highest correlation coefficient with the target variable in the training image session were selected to decode each feature (with a maximum of 500 voxels).

### Deep neural network features

We used the *Caffe* implementation (Jia et al., 2014) of the *VGG19* deep neural network (DNN) model (Simonyan & Zisserman, 2015; available from http://github.com/BVLC/caffe/wiki/Model-Zoo). All visual images were resized to 224 × 224 pixels to compute outputs by the VGG19 model. The VGG19 model consisted of a total of sixteen convolutional layers and three fully connected layers. The outputs from the units in each of these 19 layers were treated as a vector in the following decoding and reconstruction analysis. In this study, we named five groups of convolutional layers as DNN1–5 (DNN1: conv1_1, and conv1_2; DNN2: conv2_1, and conv2_2; DNN3: conv3_1, conv3_2, conv3_3, and conv3_4; DNN4: conv4_1, conv4_2, conv4_3, and conv4_4; and DNN5: conv5_1, conv5_2, conv5_3, and conv5_4), and three fully-connected layers as DNN6–8 (DNN6: fc6; DNN7: fc7; and DNN8: fc8). Basically, we used the original architecture of the VGG19 model to compute feature unit activities, but for analyses with fMRI data from the imagery experiment, we changed the DNN architecture so that max pooling layers were replaced by average pooling layers, and ReLU activation function were replaced by leaky ReLU activation function with a negative slope of 0.2 (see Simonyan & Zisserman (2015) for the details of the original DNN architecture).

### DNN feature decoding analysis

We constructed multivoxel decoders to decode the DNN feature vector of a seen image from fMRI activity patterns from training natural-image sessions (training dataset) using a set of linear regression models. In this study, we used the sparse linear regression algorithm (SLR; Bishop, 2006), which can automatically select important voxels for decoding, by introducing sparsity into weight estimation through Bayesian parameters estimation with the automatic relevance determination (ARD) prior (see Horikawa & Kamitani, 2017 for detailed description). The decoders were trained to decode the values of individual units in the feature vectors of all DNN layers using the training dataset (one decoder for one DNN feature unit), and applied to test datasets. For details of the general procedure of feature decoding, see Horikawa & Kamitani (2017).

For test datasets, fMRI samples corresponding to the same stimulus or imagery were averaged across trials to increase the signal-to-noise ratio of the fMRI signals. To compensate for a possible difference of the signal-to-noise ratio between training and test samples, the decoded features of individual DNN layers were normalized by multiplying a single scalar so that the norm of the decoded vectors of individual DNN layers matched with the mean norm of the true DNN feature vectors computed from independent 10,000 natural images. Then, this norm-corrected vector was subsequently provided to the reconstruction algorithm. See Supplementary Methods for details of the norm-correction procedure.

### Reconstruction from a single DNN layer

Given a DNN feature vector decoded from brain activity, an image was generated by solving the following optimization problem (Mahendran & Vedaldi, 2014).

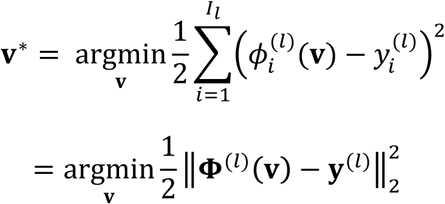
where **v** ∈ ℝ^224×224×3^ is a vector whose elements are pixel values of an image, and **v**\* is the reconstructed image. 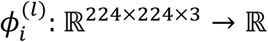 is the feature extraction function of the *i*-th DNN feature in the *l*-th layer. Namely, 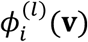 is the output value from the *i*-th DNN unit in the *l*-th layer for the image **v**. is the number of the units in the *l*-th layer. 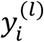 is the value decoded from brain activity for the *i*-th feature in the *l*-th layer. For simplicity, the same cost function was rewritten with a vector function in the second line. 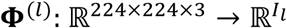 is the function whose *i*-th element is 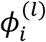 and 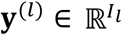 is the vector whose *i*-th element is 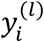.

The above cost function was minimized by limited-memory BFGS (L-BFGS; Le et al., 2011; Liu & Nocedal, 1989; Gatys et al., 2016) or by gradient descent with momentum (Qian, 1999). L-BFGS was used unless otherwise stated. The obtained solution was taken as the reconstructed image from brain activity. See Supplementary Methods for details of optimization methods.

### Reconstruction from multiple DNN layers

To combine DNN features from multiple layers, we took a weighted sum of the cost functions for individual DNN layers, which is given by

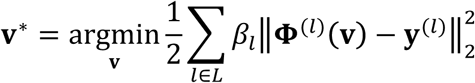
where *L* is a set of DNN layers and *β_l_* is a parameter that determines the contribution of the *l*-th layer. We set *β_l_* to 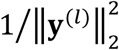 for balancing the contributions of individual DNN layers. Again, this cost function was minimized by the L-BFGS algorithm. The DNN layers included in *L* were combined. In the main analyses, we combined all convolutional layers (DNN1–5) and fully connected layers (DNN6–8) unless otherwise stated.

### Natural image prior

To improve the “naturalness” of reconstructed images, we modified the reconstruction algorithm by introducing a constraint. To constrain resultant images from all possible pixel contrast patterns, we reduced the degrees of freedom by introducing a generator network derived by the generative adversarial network algorithm (GAN; Goodfellow et al., 2014), which have been recently shown to work well to capture a latent space that explains natural images (Radford et al., 2015). In the GAN framework, a set of two neural networks, which are called a generator and a discriminator, are trained. The generator is a function to map from a latent space to the data space (i.e. pixel space), and the discriminator is a classifier that predicts whether a given image is a sample from real natural images or an output from the generator. The discriminator is trained to increase its predictive power, and the generator is trained to decrease it. We considered constraining our reconstructed image to be in the subspace that consists of the images a generator trained to produce natural images could produce (Nguyen et al., 2016; Dosovitskiy & Brox, 2016). This is expressed by

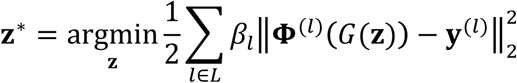
and

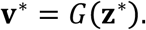

*G* is the generator as the mapping function from the latent space to the image space, which we called a deep generator network (DGN). In our reconstruction analysis, we used a pre-trained DGN which was provided by Dosovitskiy & Brox (2016; available from http://github.com/dosovits/caffe-fr-chairs; trained model for fc7).

The above cost function for the reconstruction with respect to **z** was minimized by gradient descent with momentum. We used the zero vector as the initial value. To keep **z** being within a moderate range, we restricted the range of each element of **z** following the previous study (Dosovitskiy & Brox, 2016).

### Reconstruction quality evaluation

Reconstruction quality was evaluated by either objective or subjective assessment. For the objective assessment, we performed a pairwise similarity comparison analysis, in which a reconstructed image was compared with two candidate images (its original image and a randomly selected image) and was tested whether pixel-wise spatial correlation coefficient (Pearson correlation) was higher for the original image. For the subjective assessment, we conducted a behavioral experiment with another group of 9 raters (4 females and 5 males, aged between 19 and 36 years). On each trial of the experiment, the raters viewed a display presenting a reconstructed image (at the bottom) and two candidate images (at the top; its original image and a randomly selected image), and were asked to select the image similar to the one presented at the bottom from the two candidates. Each trial continued until the raters made a response. For both types of assessments, the proportion of trials, in which the original image was selected as more similar one was calculated as a quality measure. In both objective and subjective assessments, each reconstructed image was tested with all pairs of the images among the same types of images (natural-images, geometric-shapes, and alphabetical-letters for images from the image presentation sessions, and natural-images and geometric-shapes for images from the imagery session; e.g., for the test natural-images, one of the 50 reconstructions was tested with 49 pairs consisted of one original image and another image from the rest of 49, resulting in 50 **×** 49 = 2,450 comparisons).

To compare the reconstruction quality across different combinations of DNN layers, we also conducted another behavioral experiment with another group of 4 raters (1 females and 3 males, aged between 23 and 36 years). On each trial of the experiment, the raters viewed a display presenting one original image (at the top) and two reconstructed images for the same original image obtained from different combinations of DNN layers (at the bottom), and were asked to judge which of the two reconstructed images was better. This pairwise comparison was conducted for all pairs of the combinations of DNN layers (28 pairs) and for all stimulus images presented in the test natural-image session (50 samples). Each trial continued until the raters made a response. We calculated the proportion of trials, in which the reconstructed image obtained from a specific combination of DNN layers is judged as better, then this value was treated as the winning percentage of this combination of DNN layers.

These assessments were performed for each set of reconstructions from different subjects and datasets individually (e.g., test natural-images from Subject 1). For the subjective assessments, one set of reconstructed images was tested with at least three raters. The evaluation results from different raters were averaged within the same set of reconstructions and treated in the same manner to the evaluation results from the objective assessment.

### Statistics

We used signed-rank tests to examine differences of assessed reconstruction quality from different conditions. ANOVA was used to examine interaction effects between task types and brain areas.

Supplementary Information is available in the online version of the paper.

## Acknowledgements

The authors thank Mitsuaki Tsukamoto, and Hiroaki Yamane for help with data collection, and Mohamed Abdelhack for the comments on the manuscript. This research was supported by grants from the New Energy and Industrial Technology Development Organization (NEDO), JSPS KAKENHI Grant number JP26119536, JP15H05920, JP15H05710, JP17K12771 and ImPACT Program of Council for Science, Technology and Innovation (Cabinet Office, Government of Japan).

## Author Contributions

Y.K. designed the study. S.G. and K.M. developed and implemented the algorithm. T.H. performed experiments and analyses. T.H., K.M., and Y.K. wrote the paper.

## Author Information

The authors declare no competing financial interests. Correspondence and requests for materials should be addressed to Y.K.

## Supplementary Methods

### Localizer experiments

The retinotopy and functional localizer experiments were conducted to identify the seven visual areas analyzed in the study.

The retinotopy experiments were conducted according to the conventional protocol (Engel et al., 1994; Sereno et al., 1995). We used a rotating wedge and an expanding ring covered in a flickering checkerboard. The data were used to delineate the borders between visual cortical areas, and to identify the retinotopic map (V1–V4) on the flattened cortical surfaces of individual subjects.

We also performed functional localizer experiments to identify the lateral occipital complex (LOC) (Kourtzi and Kanwisher, 2000), fusiform face area (FFA) (Kanwisher et al., 1997), and parahippocampal place area (PPA) (Epstein & Kanwisher, 1998) for each individual subject. The localizer experiment consisted of 8 runs and each run contained 16 stimulus blocks. In this experiment, intact or scrambled images (12 x 12 degrees of visual angle) from face, object, house, and scene categories were presented at the center of the display. Each of eight stimulus types (four categories x two conditions) was presented twice per run. Each stimulus block consisted of a 15-second intact or scrambled stimulus presentation. The intact and scrambled stimulus blocks were presented successively (the order of the intact and scrambled stimulus blocks was random), followed by a 15-second rest period consisting of a uniform gray background. Extra 33-second and 6-second rest periods were added to the beginning and end of each run, respectively. In each stimulus block, 20 different images of the same type were presented for 0.3 seconds, followed by an intervening blank screen of 0.4 seconds.

### MRI acquisition for localizer experiments

fMRI data were collected using 3.0-Tesla Siemens MAGNETOM Verio scanner located at the Kokoro Research Center, Kyoto University. An interleaved T2*-weighted gradient-EPI scan was performed to acquire functional images covering the entire occipital lobe (retinotopy experiment: TR, 2,000 ms; TE, 30 ms; flip angle, 80 deg; FOV, 192 × 192 mm; voxel size, 3 × 3 × 3 mm; slice gap, 0 mm; number of slices, 30) or the entire brain (localizer experiment: TR, 3,000 ms; TE, 30 ms; flip angle, 80 deg; FOV, 192 × 192 mm; voxel size, 3 × 3 × 3 mm; slice gap, 0 mm; number of slices, 46). T2-weighted turbo spin echo images were scanned to acquire high-resolution anatomical images of the same slices used for the EPI (retinotopy experiment: TR, 6,000 ms; TE, 57 ms; flip angle, 160 deg; FOV, 192 × 192 mm; voxel size, 0.75 × 0.75 × 3.0 mm; localizer experiments: TR, 7,020 ms; TE, 69 ms; flip angle, 160 deg; FOV, 192 × 192 mm; voxel size, 0.75 × 0.75 × 3.0 mm).

### MRI data preprocessing for data from localizer experiments

The first 8-second scans for experiments with TR = 2 seconds (retinotopy experiments) and 9-second scans for experiments with TR = 3 seconds (localizer experiment) of each run were discarded to avoid MRI scanner instability. We then subjected the acquired fMRI data to three-dimensional motion correction with SPM5 (http://www.fil.ion.ucl.ac.uk/spm). Those data were then coregistered to the within-session high-resolution anatomical images of the same slices used for EPI and subsequently to the whole-head high-resolution anatomical images. The coregistered data were then re-interpolated as 2 × 2 × 2 mm voxels.

### Region of interest (ROI) selection

V1, V2, V3, and V4 were identified using the data from the retinotopy experiments (Engel et al., 1994; Sereno et al., 1995). LOC, FFA, and PPA were identified using the data from the functional localizer experiments (Kanwisher et al., 1997; Kourtzi and Kanwisher, 2000; Epstein & Kanwisher, 1998). The data from the retinotopy experiment were transformed into Talairach space and the visual cortical borders were delineated on the flattened cortical surfaces using BrainVoyager QX (http://www.brainvoyager.com)(RRID: SCR_013057). The coordinates of voxels around the gray-white matter boundary in V1–V4 were identified and transformed back into the original coordinates of the EPI images. The localizer experiment data were analyzed using SPM5. The voxels showing significantly higher activation in response to intact object, face, or scene images compared with that for scrambled images (t-test, uncorrected *p* < 0.05 or 0.01) were identified, and defined as LOC, FFA, and PPA respectively. A contiguous region covering LOC, FFA, and PPA was manually delineated on the flattened cortical surfaces, and the region was defined as the “higher visual cortex” (HVC). Voxels from V1–V4 and the HVC were combined to define the “visual cortex” (VC). In the regression analysis, voxels showing the highest correlation coefficient with the target variable in the training image session were selected to decode each feature (with a maximum of 500 voxels).

### Norm correction for decoded DNN feature vectors

Before reconstruction analysis, the DNN feature vector decoded from a given fMRI sample was multiplied by a scalar to match its norm to the mean across natural images.

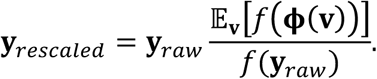
**y***_raw_* is a decoded feature vector, and **y***_rescaled_* is the feature vector after the norm-correction. **v** is a vector whose elements are pixel values of an image, and **ϕ** is the feature extraction function whose input is an image vector and output is the DNN feature vector for the input image. *f* is a function whose definition is given later, and 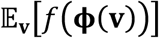 denotes the expectation of 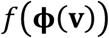 with respect to **v** across natural images. The expectation was calculated using 10,000 natural images randomly selected from the *ImageNet* database (Deng et al., 2009; 2011, fall release).

When the input of the function *f* is a DNN feature vector from a convolutional layer, we first calculate the standard deviation of the feature value across the units in each channel, then the mean of this standard deviation across all channels is treated as the output value of *f*. When the input is a DNN feature vector from a fully-connected layer, the standard deviation of the feature value across the units in the layer is treated as the output value of *f*.

If *f* is the vector norm, our norm-correction exactly matches the given decoded vector with the mean norm across natural images. In this study, we adopted the definition of *f* explained above because our norm-correction procedure led to slightly better reconstructions compared to the exact norm matching in analysis at early stages with independent preliminary data.

### Optimization methods for reconstruction

The cost function for our reconstruction was minimized by limited-memory BFGS (L-BFGS; Le et al., 2011; Liu & Nocedal, 1989; Gatys et al., 2016) or by gradient descent with momentum (Qian, 1999). Each of those algorithms was explained in this section.

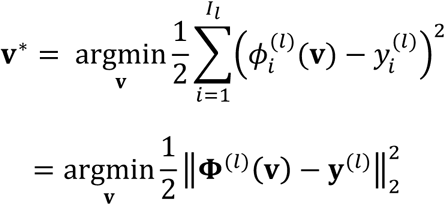
where **v** ∈ ℝ^224×224×3^ is a vector whose elements are pixel values of an image, and **v\*** is the reconstructed image. 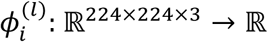 is the feature extraction function of the *i*-th CNN feature in the *l*-th layer. Namely, 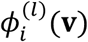 is the output value from the *i*-th DNN unit in the *l*-th layer for the image **v**. *I_l_* is the number of the units in the *l*-th layer. 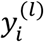 is the value decoded from brain activity for the *i*-th feature in the *l*-th layer. For simplicity, the same cost function was rewritten with a vector function in the second line. 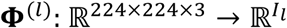 is the function whose *i*-th element is 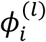 and 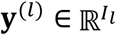 is the vector whose *i*-th element is 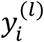.

In each iteration of the L-BFGS algorithm, the image was updated by

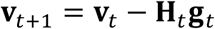
where **v***_t_* and **v***_t_*_+1_ are the vectors before and after the *t*-th update. **g***_t_* is the gradient of the cost function at **v***_t_*. **H***_t_* is an approximation of the inverse hessian of the cost function at **v***_t_*.

For each update, this gradient was calculated by the backpropagation algorithm as follows. Here, we define the backpropagated error 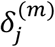 by

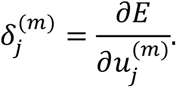
where *E* is the cost function to be minimized and 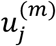 is the input to the *j*-th unit in the *m*-th layer in the forward path. Using the chain rule, 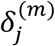 can be calculated as a weighted sum of the backpropagated errors for the units in the (*m* + 1)-th layer.

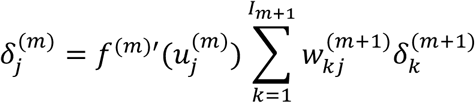
where the function *f*^(^*^m^*^)^′ is the derivative of the activation function between the *m*-th layer and (*m* + 1)-th layer. 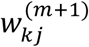 is the weight between the *j*-th unit in the *m*-th layer and the *k*-th unit in the (*m* + 1)-th layer.

The backpropagated error for a unit in last layer is given by

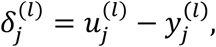
and the gradient **g***_t_* is obtained as 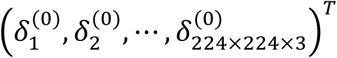, which can be numerically calculated using the chain rule.

The calculation of the inverse hessian with a size of *I_l_* × *I_l_* is intractable because it requires huge memory. To avoid the memory problem, the inverse hessian was approximated based on the history of **g***_t_* and **v***_t_* following the update rule of the L-BFGS algorithm (Liu & Nocedal, 1989). Each image was generated by 200 iterations and the spatially uniform image with the mean RGB contrast values of natural images was used as the initial image.

Also, the cost function was minimized by gradient descent with momentum (Qian, 1999). In each iteration of the algorithm, the image was updated by

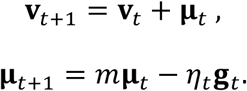
**v***_t_* and **v***_t_*_+1_ are the vectors before and after the *t*-th update. **g***_t_* is the gradient of the cost function at **v***_t_*, and **μ***_t_* is a weighted average of the gradients from step 0 to *t*. The next update is determined based on the history of **g***_t_* to prevent **v***_t_* from oscillating around shallow local minima of the cost function. *m* is a parameter which is called the decay rate, and we set this to 0.9. *η_t_* is the learning rate. Each image was generated by 200 iterations and *η_t_* was linearly reduced from 2.0 to 0.0. As the initial image for optimization, the spatially uniform image with the mean RGB contrast values of natural images was used.

For each update, the gradient **g***_t_* was calculated by the backpropagation algorithm with the procedure same as the L-BFGS algorithm.

**Supplementary Figure 1.**
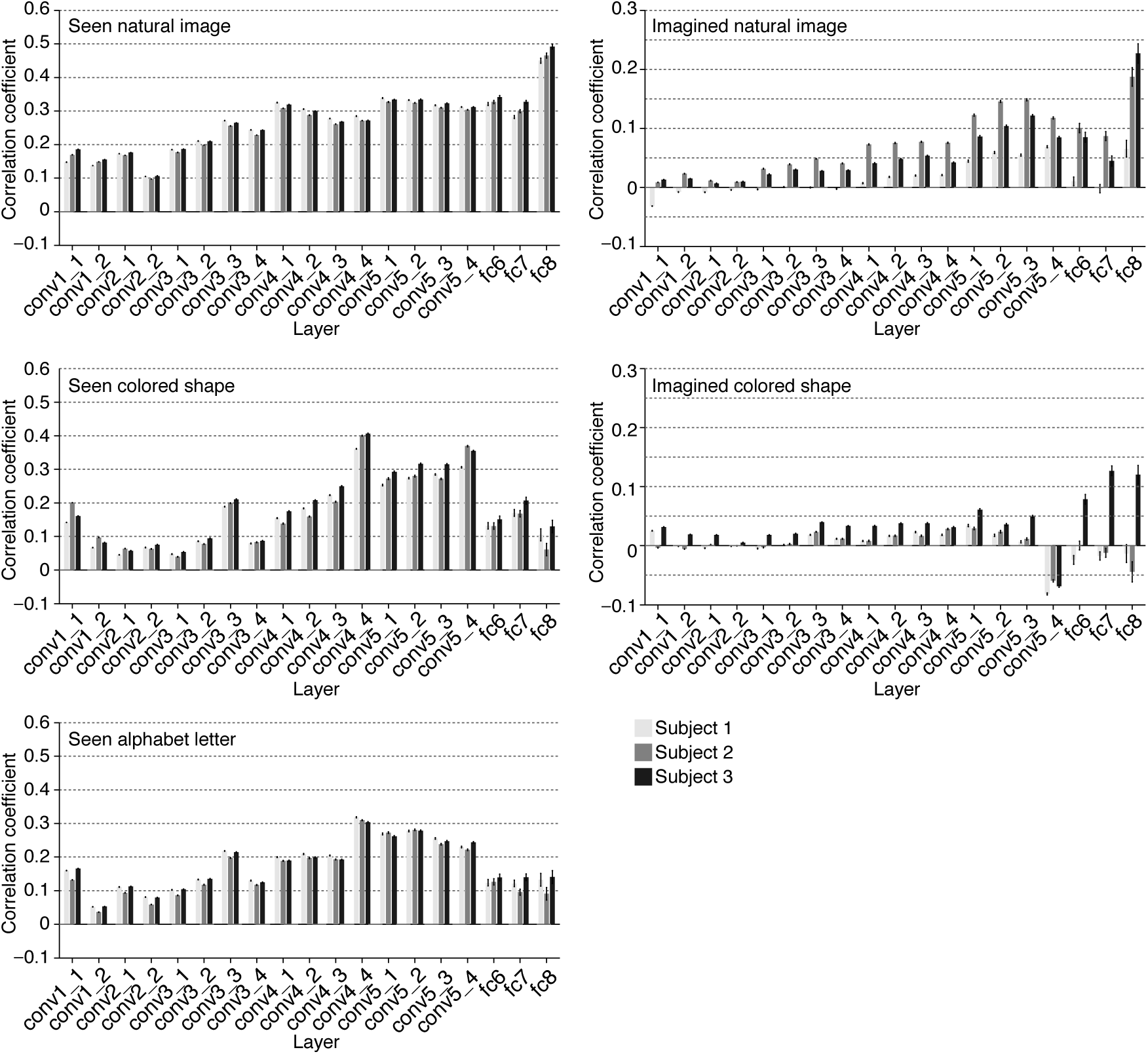
DNN feature decoding accuracy. DNN feature decoding accuracy obtained from VC activity was evaluated by the correlation coefficient between true and decoded feature values of each feature unit following the same procedure in Horikawa & Kamitani (2017). The evaluation was individually performed for each of three types of seen images (natural image, colored shapes, and alphabet letter) and each of two types imagery images (natural image and colored shape). Correlation coefficients were averaged across units in each DNN layer. Mean correlation coefficients are shown for each types of layers (error bars, 95% CI across units).

**Supplementary Figure 2.**
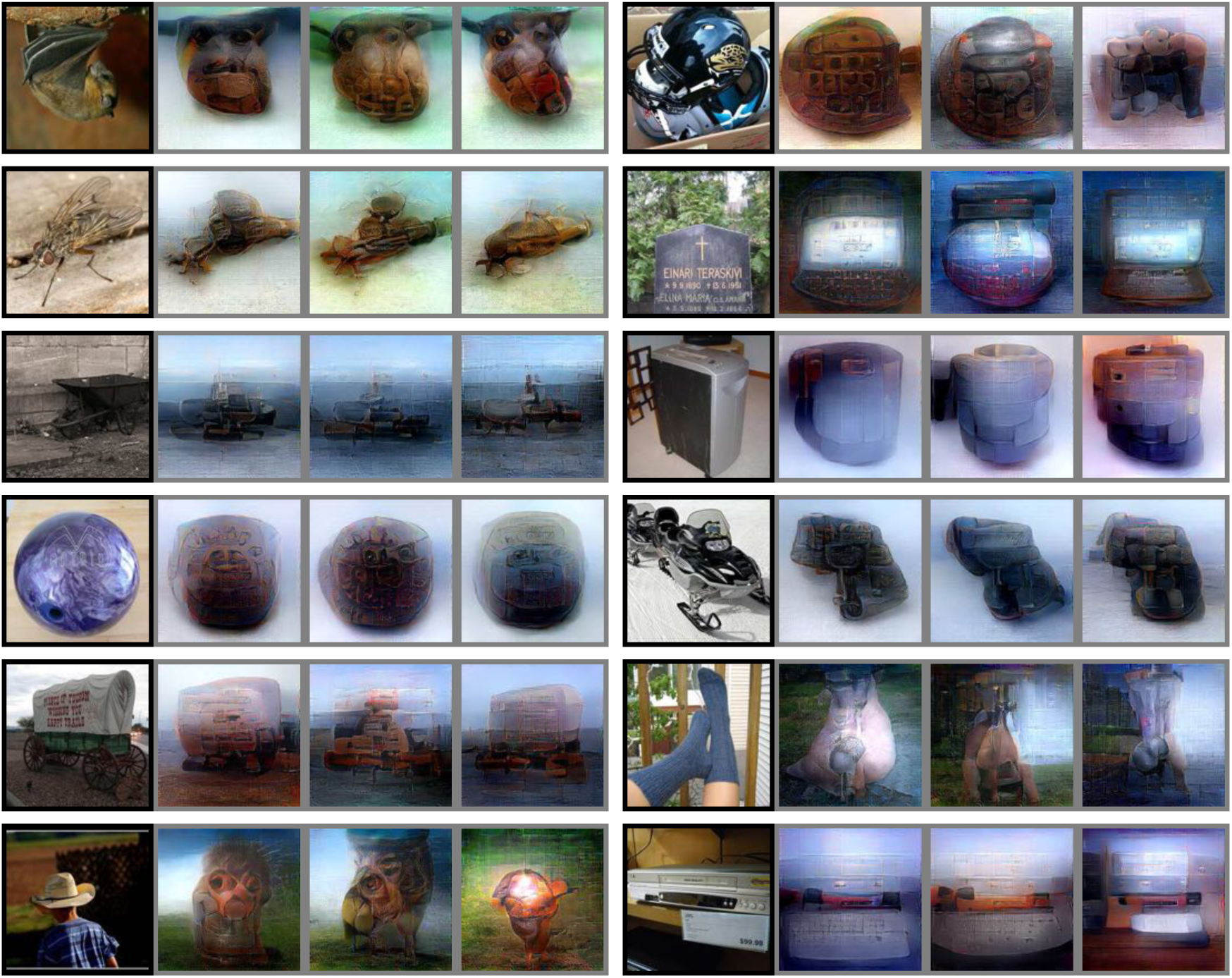
Other examples of natural image reconstructions obtained with the DGN. Images with black; and gray frames show presented and reconstructed images, respectively (reconstructed from VC activity using all DNN layers). Three reconstructed images correspond to reconstructions from three subjects.

**Supplementary Figure 3.**
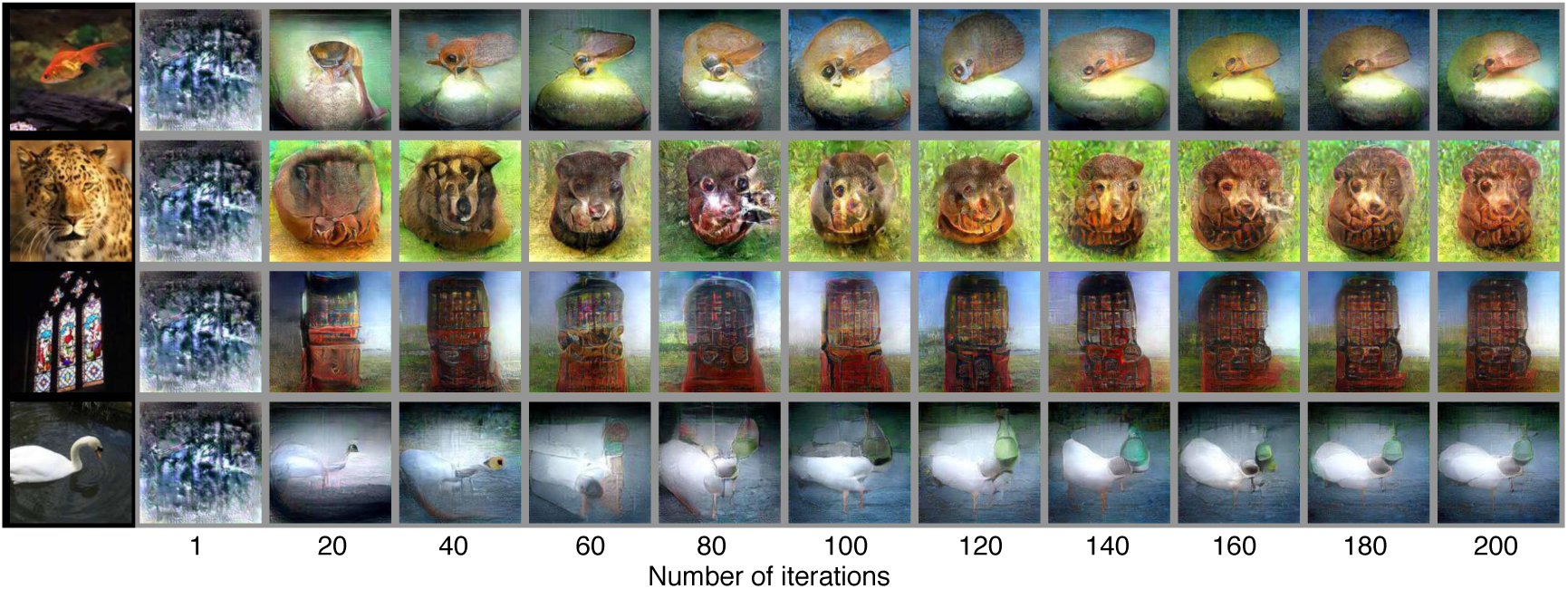
Reconstructions through optimization processes. Reconstructed images obtained through the optimization processes are shown (reconstructed from VC activity of Subject 1 using all DNN layers and the DGN). Images with black and gray frames show presented and reconstructed images, respectively.

**Supplementary Figure 4.**
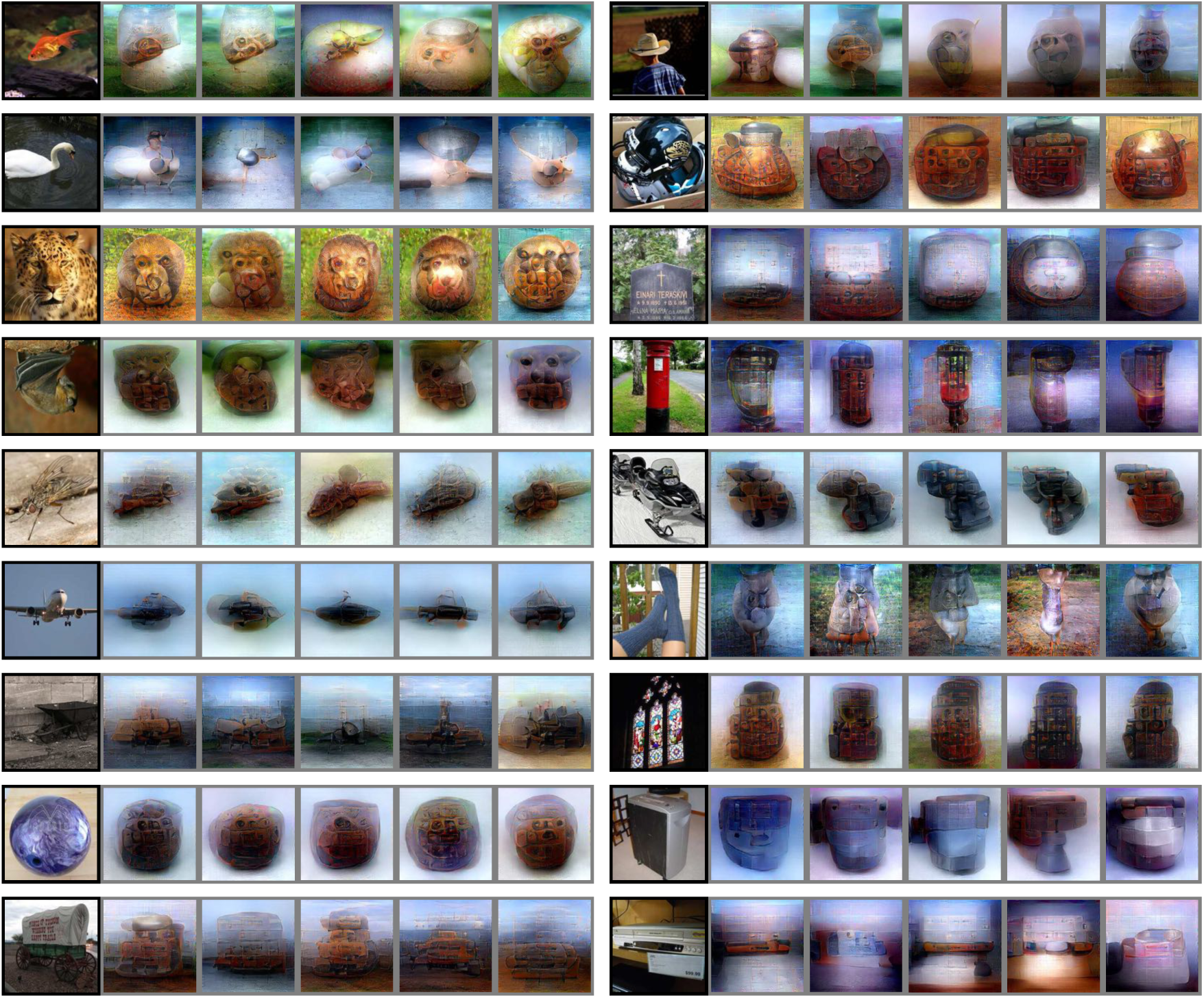
Reconstructions from the generic object decoding dataset. The same reconstruction analysis was performed with a previously published dataset (Horikawa & Kamitaai, 2017; reconstructed from VC activity using all DNN layers and the DGN). See Horikawa & Kamitani 2017; for details of tine data. Images with black and gray? frames show presented and reconstructed images, respectively. Five reconstructed images correspond to reconstructions from five subjects.

**Supplementary Figure 5.**
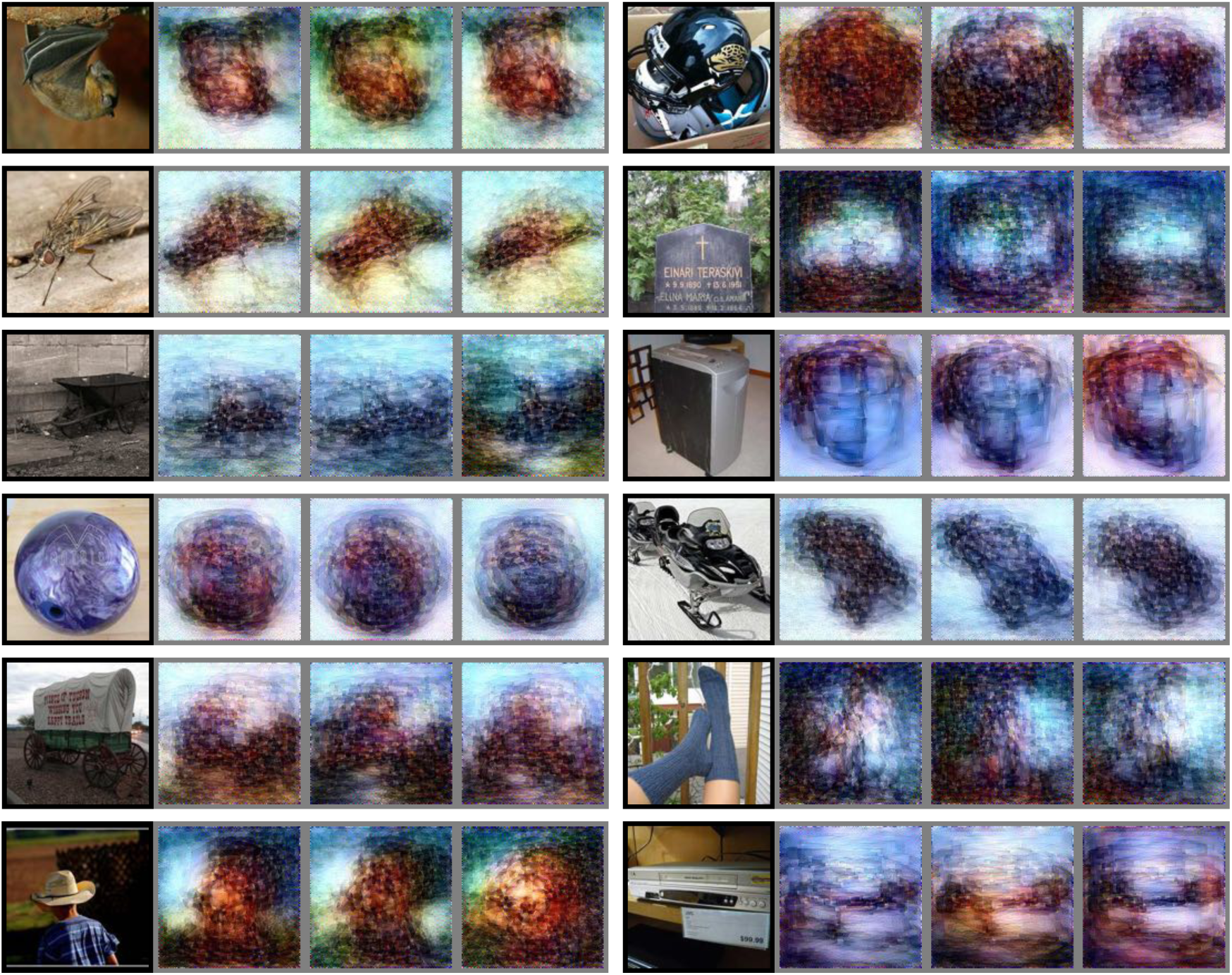
Other examples of natural image reconstructions obtained without the DGN. Images with black and gray frames show presented and reconstructed images, respectively (reconstructed from VC activity using all DNN layers). Three reconstructed images correspond to reconstructions from three subjects.

**Supplementary Figure 6.**
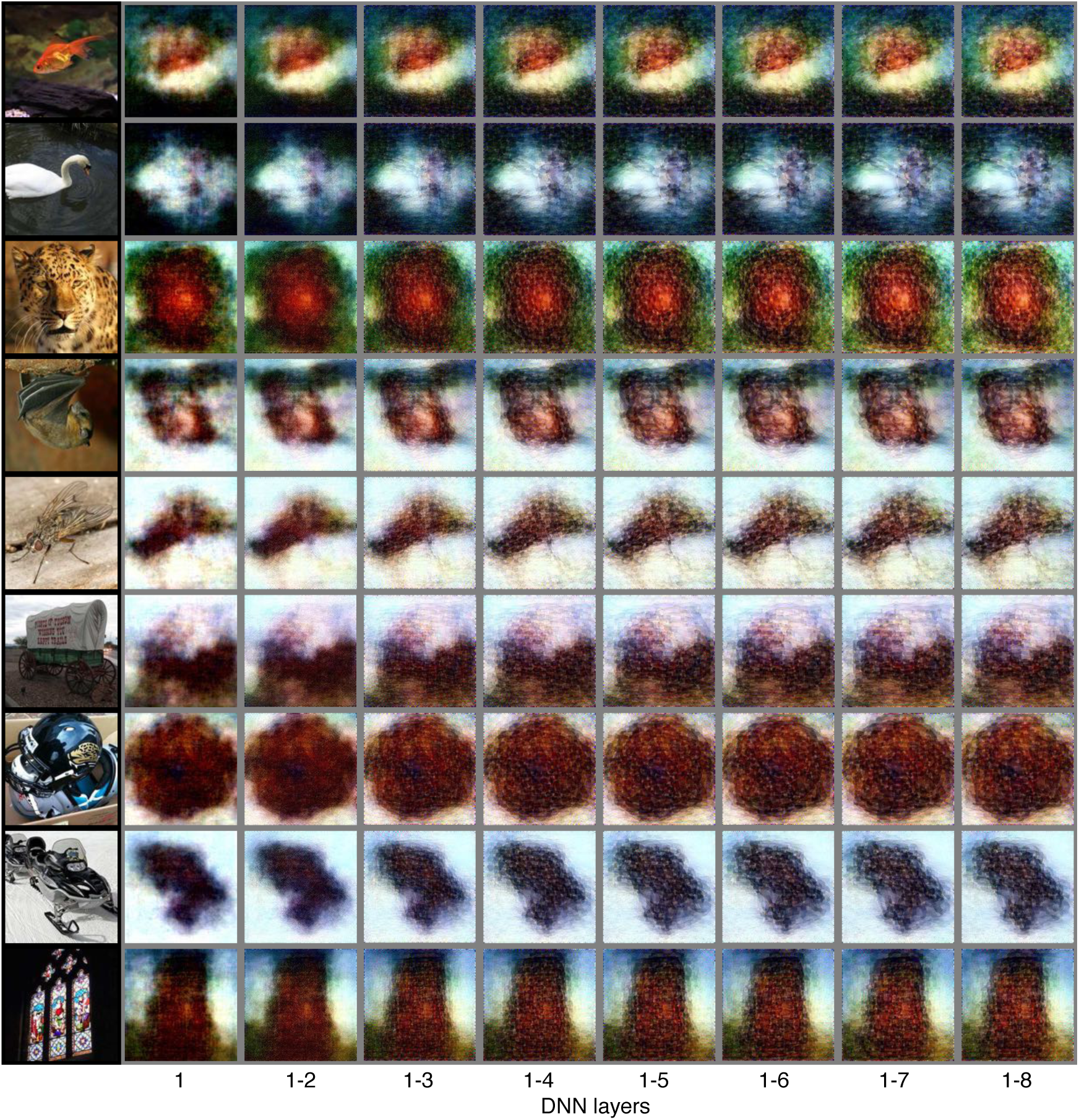
Reconstructions using different combinations of DNN layers (without the DGN). Images with black and gray frames show presented and reconstructed images, respectively (reconstructed from VC activity).

**Supplementary Figure 7.**
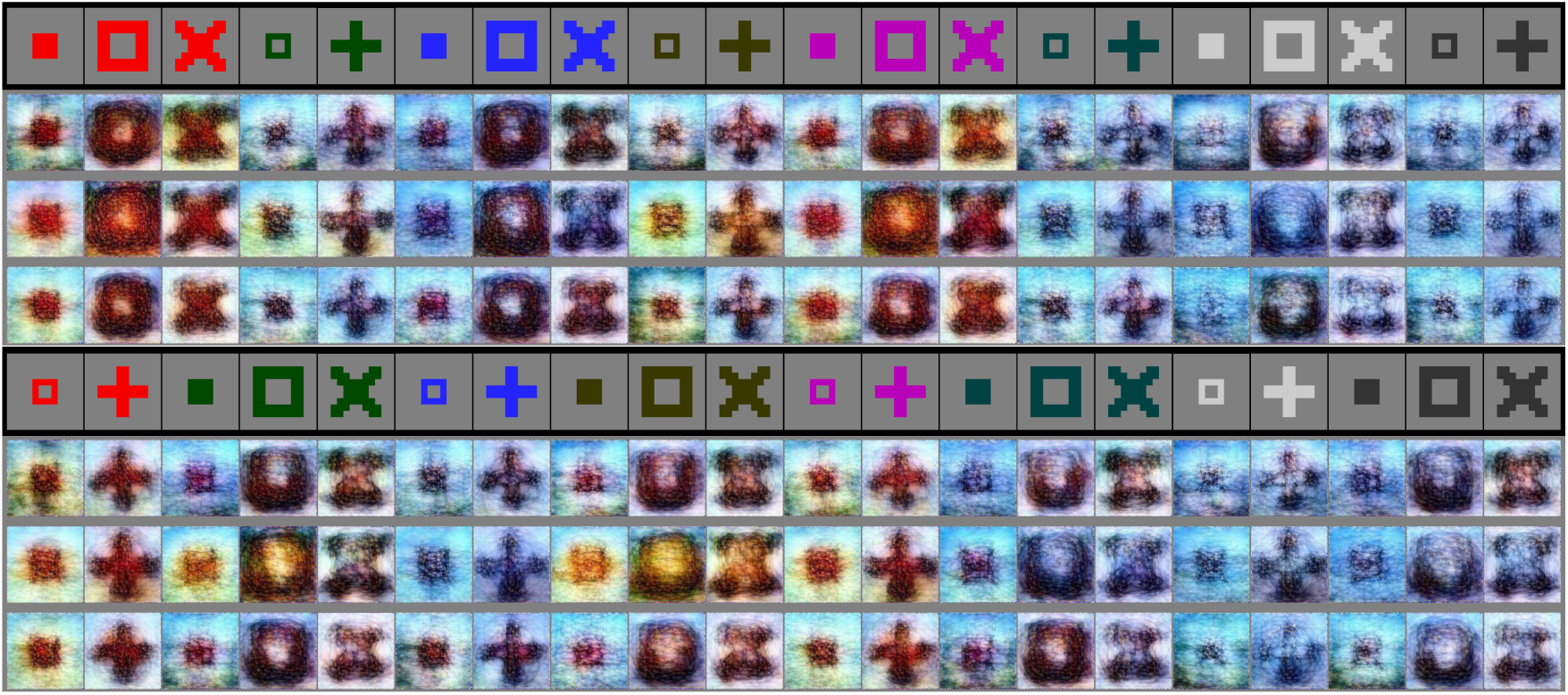
All examples of artificial colored shape reconstructions. Images with black and gray frames show presented and reconstructed images, respectively (reconstructed from VC activity). Three reconstructed images correspond to reconstructions from three subjects.

**Supplementary Figure 8.**
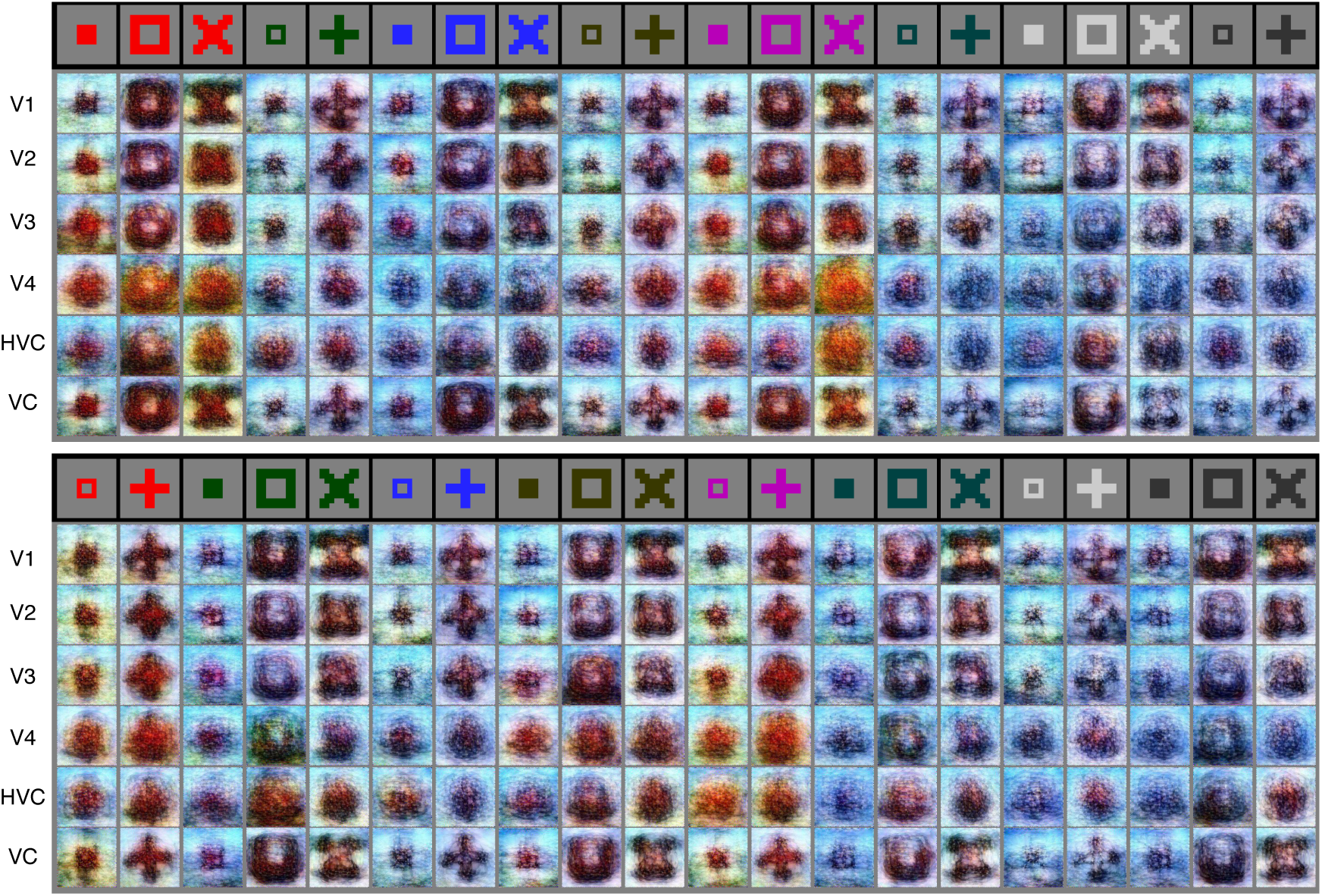
All examples of artificial colored shapes reconstructions obtained from different visual areas (Subject 1). Images with black and gray frames show presented and reconstructed images, respectively (reconstructed from VC activity using all DNN layers without the DGN).

**Supplementary Figure 9.**
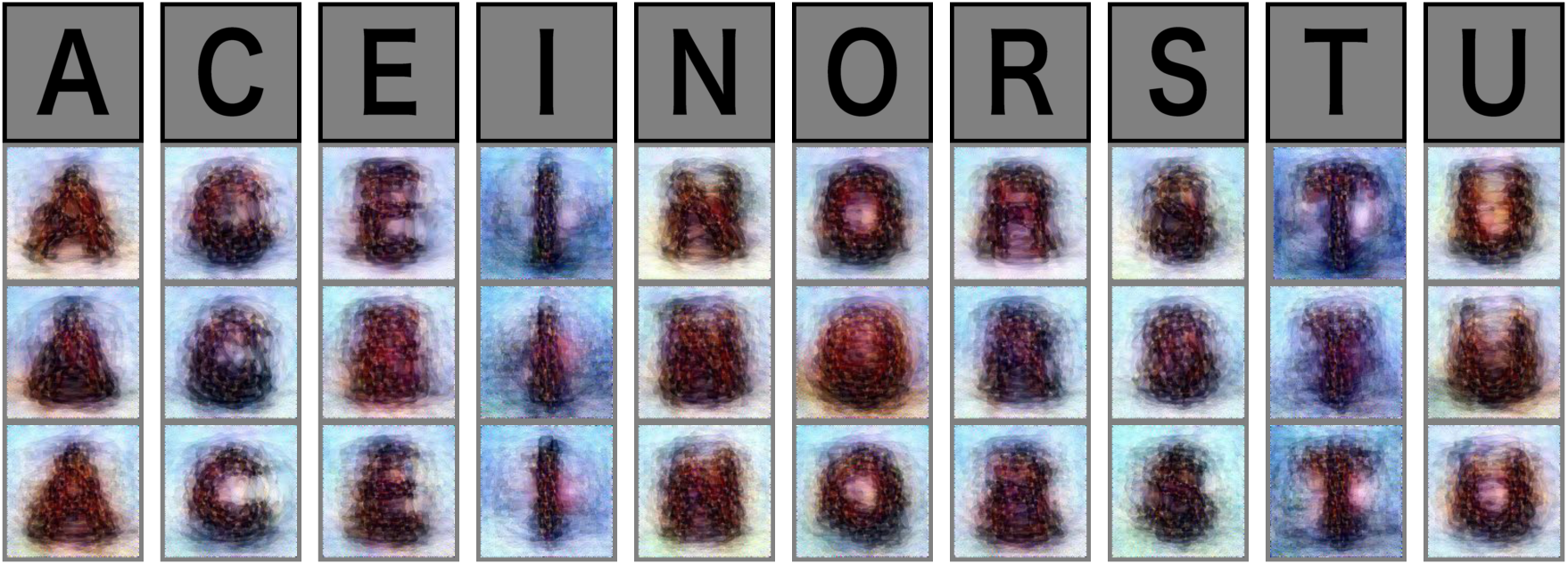
All examples of alphabetical letter reconstructions. Images with black and gray frames show presented and reconstructed images, respectively (reconstructed from VC activity without the DGN). Three reconstructed images correspond to reconstructions from three subjects.

**Supplementary Figure 10.**
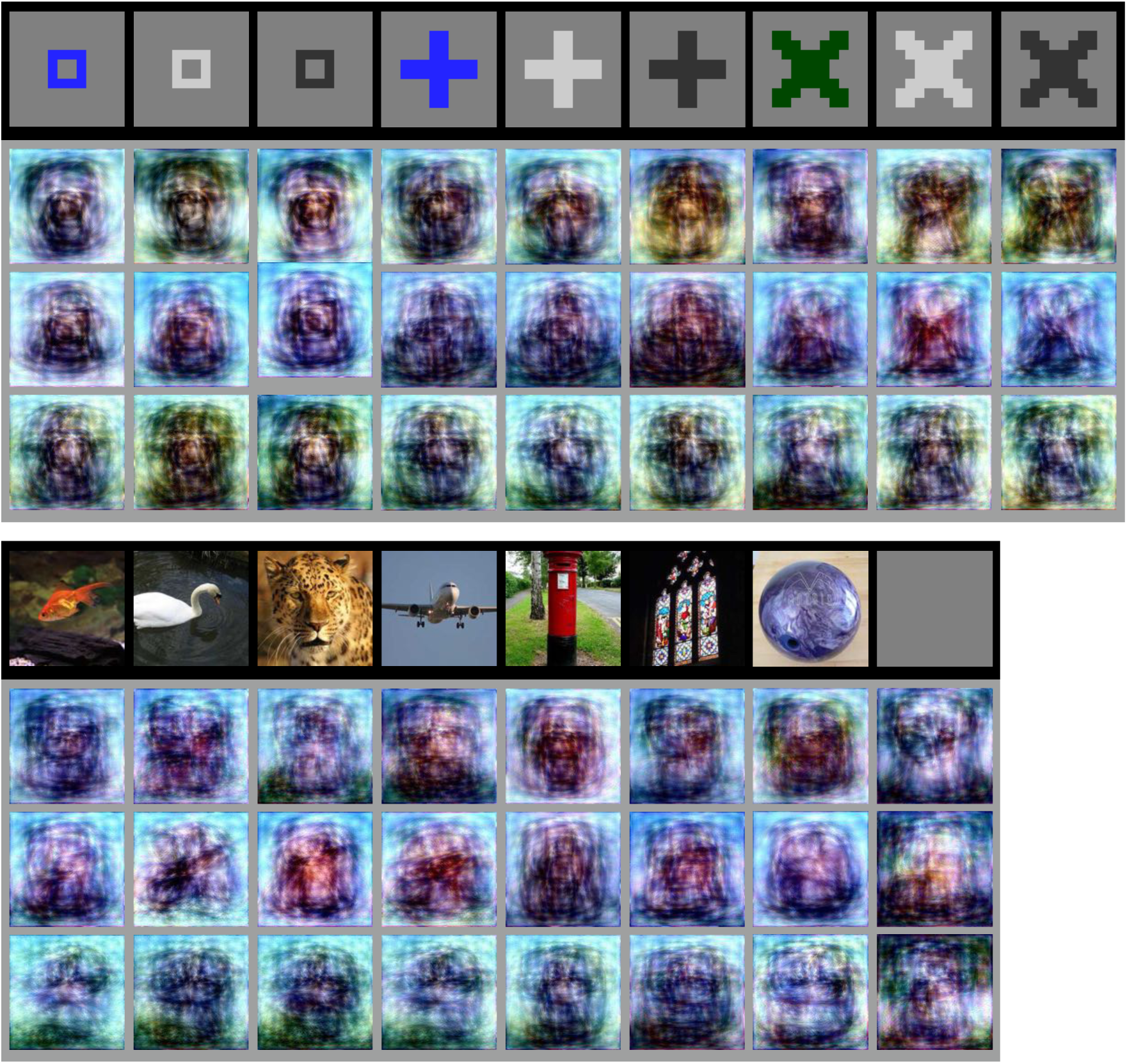
Other examples of imagery image reconstructions. Images with black and gray frames show instructed and reconstructed images, respectively (reconstructed from VC activity without the DGN). Three reconstructed images correspond to reconstructions from three subjects. Rightmost images in the bottom row show reconstructions during maintaining fixation without imagery.

**Supplementary Figure 11.**
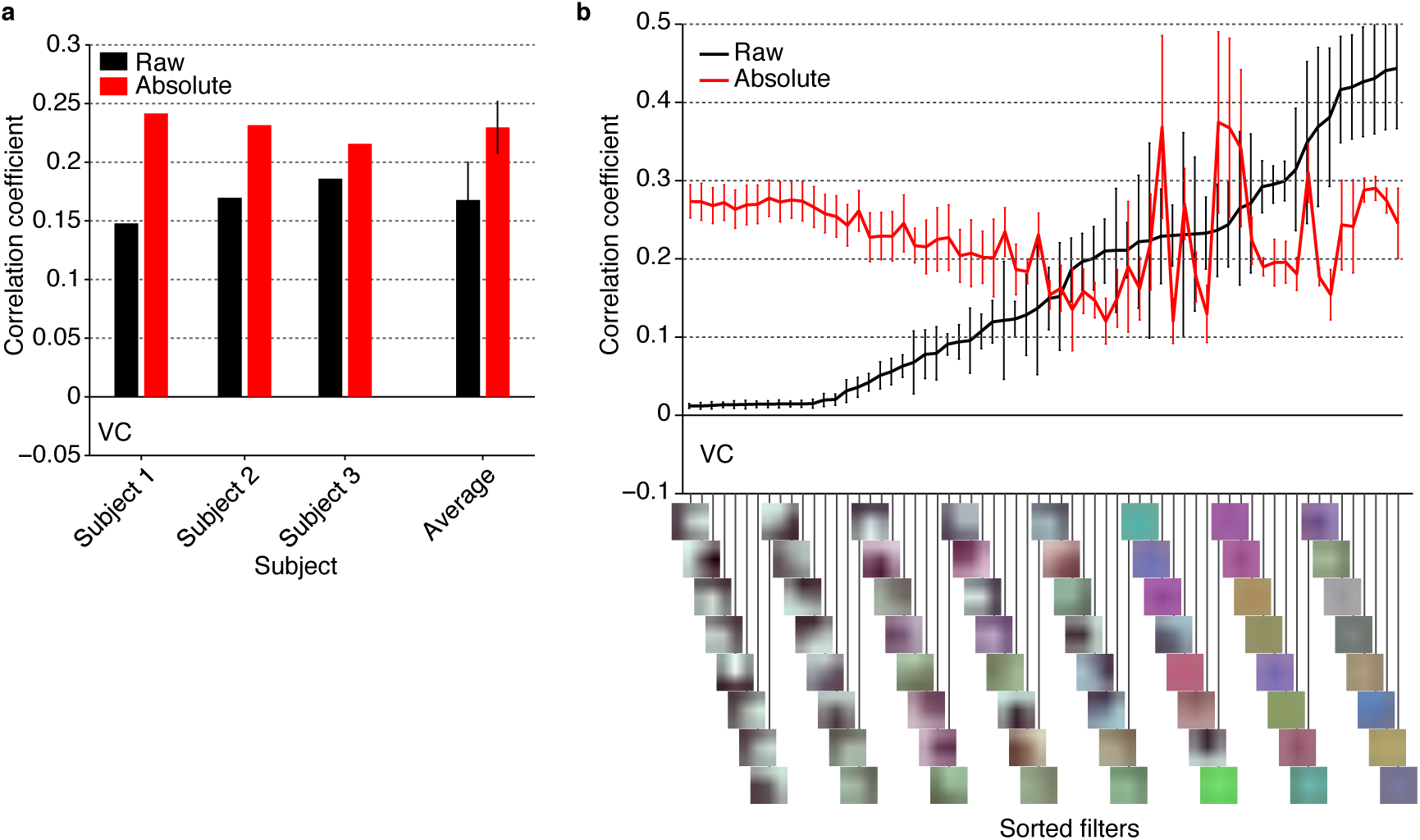
Feature decoding accuracy of raw and absolute features. In addition to the feat decoding analysis with raw DNN feature, we performed the same feature decoding analysis after converting raw feature outputs to absolute values. The analysis was performed with features from the conv1_1 layer of the VGG19 model using the natural object dataset (error bar, 95%) C.I. across subjects). a, Mean feature decoding accuracy of all units. The results showed significant improvements of feature decoding accuracy by the absolute conversion. b, Mean feature decoding accuracy for individual filters. The feature decoding accuracies of units within the same filters were individually averaged. The filters were sorted according to the ascending order of the raw feature decoding accuracy averaged for individual filters. These results showed that feature decoding accuracies of monochrome colored filters were specifically improved by the conversion. The large improvement levels demonstrate the insensitivity of fMRI signals to pixel luminance, suggesting the linear-nonlinean discrepancy of DNN and fMRI responses to pixel luminance. This discrepancy may? explain the reversal of luminance observed in several reconstructed images.

## Notes for Supplementary Movies

Reconstruction of visual images from human brain activity measured by fMRI. To reconstruct visual images, we first decoded (translated) measured brain activity patterns into deep neural network (DNN) features, then fed those decoded features to a reconstruction algorithm. Our reconstruction algorithm starts from a given initial image and iteratively optimizes the pixel values so that the DNN features of the current image become similar to those decoded from brain activity. The movies can be seen from our repository: http://www.youtube.com/user/ATRDNI

Movie. Deep image reconstruction: Natural images.

## Footnotes

1 http://www.youtube.com/user/ATRDNI

